# Comprehensive Virulence Profiling and Evolutionary Analysis of Specificity Determinants in *Staphylococcus aureus* Two-Component Systems

**DOI:** 10.1101/2023.10.20.563342

**Authors:** Stephen Dela Ahator, Karoline Wenzl, Kristin Hegstad, Christian Lentz, Mona Johannessen

## Abstract

In the *Staphylococcus aureus* genome, a set of highly conserved two-component systems (TCSs) composed of histidine kinases (HKs) with their cognate response regulators (RRs) sense and respond to environmental stimuli, which drive the adaptation of the bacteria. This study investigates the complex interplay between TCSs in *S. aureus* USA300, a predominant Methicillin-Resistant *S. aureus* (MRSA) strain, revealing shared and unique virulence regulatory pathways and genetic variations mediating signal specificity within TCSs. Using TCS-related mutants from the Nebraska Transposon Mutant Library, we analyzed the effects of inactivated TCS HKs and RRs on the production of various virulence factors, *in vitro* infection abilities, and adhesion assays. We found that the TCSs influence on virulence determinants was not associated with their phylogenetic relationship, indicating divergent functional evolution. Using the cocrystalized structure of the DesK-DesR from *B. subtilis* and modelled structures of the 4 NarL TCSs in *S. aureus*, we identified interacting residues, revealing specificity determinants and conservation within the same TCS, even from different strain backgrounds. The interacting residues were highly conserved within strains but varied between species due to selection pressures and coevolution of cognate pairs. This study unveils the complex interplay and divergent functional evolution of TCSs, highlighting their potential for future experimental exploration of phosphotransfer between cognate and non-cognate recombinant HK and RRs.

**IMPORTANCE:** Given the widespread conservation of Two-Component Systems (TCSs) in bacteria and their pivotal role in regulating metabolic and virulence pathways, they present a compelling target for anti-microbial agents—especially in the face of rising multi-drug resistant infections. Harnessing TCSs therapeutically necessitates a profound understanding of their evolutionary trajectory in signal transduction, as this underlies their unique or shared virulence regulatory pathways. Such insights are critical for effectively targeting TCS components, ensuring an optimized impact on bacterial virulence and mitigating the risk of resistance emergence via the evolution of alternative pathways. Our research offers an in-depth exploration of virulence determinants controlled by TCSs in S. aureus, shedding light on the evolving specificity determinants that orchestrate interactions between their cognate pairs.

## INTRODUCTION

*Staphylococcus aureus*, a commensal yet opportunistic bacterium, colonizes the skin and mucous membranes of nearly 30% of healthy adults. This adaptable pathogen is responsible for various hospital and community-acquired infections due to its production of a diverse array of virulence factors. These factors include cell wall anchored proteins such as protein A, fibronectin-binding proteins, and clumping factors (1–3) and secreted exoproteins such as nucleases, proteases, lipases, collagenases, and hemolysins (1).

Bacteria have evolved sophisticated regulatory mechanisms, such as the two-component system (TCS), which integrate virulence regulation with responses to host and environmental factors (4). A typical TCS comprises a membrane-spanning sensor histidine kinase (HK) that detects stimuli, autophosphorylate, and subsequently phosphorylates a conserved aspartate residue on the cytoplasmic response regulator (RR). This second phosphorylation event induces conformational changes in the RR, enabling specific binding to DNA motifs to enact appropriate responses. The HK typically has two conserved domains: the dimerization and histidine phosphotransfer (DHp) domain for autophosphorylation and phosphotransfer and the catalytic and ATP binding (CA) domains (4, 5). HKs usually have an N-terminal transmembrane domain for cell membrane attachment and signal perception. Additional domains, such as PAS (Per Arnt Sim), HAMP (Histidine kinase, adenyl cyclases, methyl-accepting proteins and phosphatases), and GAF (cGMP-specific phosphodiesterases, adenylyl cyclases and FhlA) (6), help relay signals from sensory domains to DHp and CA (6–8). RRs contain a conserved receiver domain that facilitates phosphotransfer from the cognate HK, activating the DNA-binding output domain for transcriptional regulation (9). The TCS’s fundamental architecture is highly adaptable, with variations emerging from its modular architecture, domain shuffling, and sensor domain diversification, allowing the detection of diverse stimuli (10–13).

TCSs play a significant role in bacterial evolution, with their functional importance influencing their evolutionary history. Acquired through gene duplication and horizontal gene transfer, the maintenance of these genes depends on their short-term selective advantage or functional specialization (14, 15). Coevolution of HKs and cognate RRs occurs through mutations, ensuring specificity while avoiding unwanted cross-talk (8, 16–18). Most *S. aureus* strains contain 15-17 TCSs, classified into four families: NarL, AraC, OmpR, and LytTR, based on the homology of the RR effector domain and sequences surrounding the active-site histidine sequences of the HK (5, 19, 20). Despite specialized functionality, many TCSs in *S. aureus* have overlapping regulons, which is linked to their regulation of alternative regulatory pathways and the maximal expression of virulence determinants (8).

TCSs have been proposed as potential targets for antibacterial agents due to their functionality, conservation, and distribution (21). Although previous studies have analyzed the functions of individual TCSs independently, a comprehensive understanding of their collective influence on virulence and the determinants of specificity remains elusive. Here, we report the results of the systemic analysis of TCSs in the *S. aureus* USA300 genome, a predominant methicillin-resistant *S. aureus* (MRSA), and a major cause of severe community-acquired infections and antibiotic resistance. Using a comprehensive panel of strains from the Nebraska Transposon Mutant Library deficient in individual TCS components, we examined how TCSs control a panel of virulence determinants common to *S. aureus* and clustered them accordingly. Furthermore, the evolutionary stability of the TCSs from genomes of *S. aureus* was determined to verify the conservation and site-specific selection pressure on the components of all the TCSs in *S. aureus* USA300. Using the NarL family of TCSs and the cocrystal structures of the DesK-DesR from *B. subtilis,* we show the variations and conservation in amino acids mediating the interactions between the DHp and rec domains and how they influence the specificity of phosphotransfer in the TCSs.

## MATERIALS AND METHODS

### Bacterial strains

The *Staphylococcus aureus* strain USA300 JE2 was used as the wildtype (WT). The transposon mutants of the TCS components were obtained from the Nebraska Transposon Mutant Library (22)(Table S1). Transposon mutants were chosen based on transposon insertions impacting the gene, specifically those located within the first 60% of the gene length.

### Human cell lines

The immortalized human keratinocyte cell line HaCaT was cultured in Dulbecco’s Modified Eagle’s medium (DMEM) supplemented with 1% penicillin/streptomycin (Sigma) and 10% Fetal Bovine Serum (FBS). The human macrophage cell line THP-1 was cultured in RPMI 1640 media supplemented with 1% penicillin/streptomycin and 10% FBS. Prior to the infection assay, the THP-1 cells were induced to differentiate by adding 10 ng/ml of Phorbol myristate acetate (PMA) for 24 hours. All the cell lines were cultured at 37 °C in a 5% CO_2_ incubator.

### Protease assay

The protease assay was performed using the plate diffusion method. Overnight cultures of the *S. aureus* mutants and WT grown in TSB at 37°C were sub-cultured in fresh TSB to an OD600 of 0.5. An aliquot of 10 µl of the OD-adjusted cultures was spot inoculated onto 1.5% w/v Mueller Hinton (MH) supplemented with 2% skim milk (Sigma). Plates were incubated at 25°C for 24 hours. The zone of clearance around the colonies, which is directly proportional to the level of protease activity, was measured.

### HaCaT cell adhesion assay

The HaCaT cell lines were used to test the adhesion ability of the WT and various TCS mutants. The HaCaT cells were seeded in 24 well plates at a density of 1 ×10^6^ cells per ml in DMEM (supplemented with 10% v/v FBS, 1% Penicillin/Streptomycin. Overnight cultures of *S aureus* strains in TSB were sub-cultured to OD600 of 1 and diluted in DMEM (supplemented with 10% v/v FBS) to a density of 1 ×10^8^ CFU/ml. The HaCaT cells were added bacteria at a MOI of 10 and incubated for 30 minutes at 37°C and 5% CO_2_. The media were removed, and the cells were washed three times with sterile PBS. The cells were lysed with 0.2% triton X-100 (Sigma) in PBS. The lysed cell suspension was serially diluted in PBS, and the number of adherent bacteria was counted by plating and incubating on MH agar for 24 hours at 37°C.

### HaCaT cell viability assay

The test for HaCaT cell viability following the addition of *S. aureus* strains was performed with the tetrazolium dye, MTT (3-[4,5-dimethylthiazol-2-yl]-2,5 diphenyl tetrazolium bromide) assay which determines mitochondrial activity. The HaCaT cells were seeded at 5000 cells per well into a 96-well plate. Overnight cultures of *S. aureus* strains grown in TSB at 37°C were sub-cultured into fresh TSB and grown to OD600 of 1 and diluted to 1 ×10^7^ CFU/mL in DMEM. The HaCaT cells were added to *S. aureus* at a MOI of 10 and incubated for 1 hour. Thereafter, the cells were washed with PBS, and tetrazolium dye was added at a final concentration of 5 mg/ml. The plates were incubated for 2 hours at 37°C and 5% CO_2_. The media were aspirated from each well and the purple formazan crystals dissolved with dimethyl sulfoxide (DMSO) for 1 hour at 37°C. The absorbance proportional to the viability of the HaCaT cells was measured at 570 nm.

### THP-1 cell infection assay

The differentiated THP-1 cell lines were seeded into 24-well plates at a density of 1 ×10^5^ cells per well in RPMI-1640 (supplemented with 10% v/v FBS, 1% Penicillin/Streptomycin and 0.05 mM 2-mercaptoethanol) (Sigma) and incubated for 24 hours in 37°C and 5% CO_2_. Overnight cultures of the *S. aureus* strains in Tryptic Soy broth (TSB) were sub-cultured into fresh TSB and grown to OD600 of 1. The bacteria density was adjusted to 1×10^7^ CFU/ml in RPMI-1640 (without Penicillin/Streptomycin) and added to each well of seeded THP-1 cells at a MOI of 10. The cells were infected for 45 min, washed with PBS and incubated for 30 min in RPMI-1640 (supplemented with 100 μg/mL gentamicin and 10% FBS) to kill extracellular bacteria. The cells were washed twice with PBS and lysed with PBS containing 0.2 % v/v triton X -100 in PBS. The lysed cells were serially diluted in PBS and the number of viable bacteria was counted by plating on Mueller Hinton Agar plates for 24 hours at 37°C.

### Nuclease assay

The nuclease assay was performed using the plate diffusion method with the DNase agar (Oxoid). Overnight cultures of the *S. aureus* mutants and WT grown in TSB at 37°C were sub-cultured in fresh TSB to an OD600 of 0.5. An aliquot of 10 µl of the OD-adjusted cultures was spot-inoculated onto DNase agar and incubated at 37°C for 24 hours. To develop a better contrast between the zone of DNase activity around the colonies, the agar plates were flooded with 1N Hydrochloric acid (HCl) and allowed to penetrate the media surface for 5 min. The clear zone of polymerized DNA around the colonies indicative of DNase activity was measured.

### Biofilm assay

Overnight cultures of *S. aureus* strains grown in TSB were sub-cultured on fresh TSB and grown to OD600 of 0.5. Briefly, 1 μL of the culture was transferred to 200 μL of TSB in 96-well plates and incubated overnight at 37°C without agitation. The wells were washed to remove the planktonic cells and the plates were air-dried before staining with 200 μL of 0.1% w/v Crystal violet for 10 min at room temperature. The wells were washed to remove excess dye and blotted to dry. Each well was filled with 200 μL of DMSO and incubated for 10 min at room temperature to dissolve the dye. The absorbance was measured at 590 nm.

### Hemolysis

The hemolysis assay was performed using the plate diffusion method. A volume of 10 μL of *S. aureus* strains grown in TSB to OD600 of 0.5 were spot inoculated onto Blood Agar plates and incubated at 37°C for 24 hours and an additional 4°C incubation for 24 hours. The diameter of the hemolytic zone was measured after 48 hours.

### Protein A assay

Costar 96-well ELISA plates were coated with 10 μg of anti-protein A antibodies (ACRIS) in PBS overnight at 4°C, washed 3 times with PBS-T (100 μL PBS containing 0.05%v/v Tween) and air-dried. Additionally, the plates were blocked with 1% Bovine Serum Albumin (BSA) for 2 hours at 37°C.

Overnight cultures of *S. aureus* strains grown in TSB at 37°C and 200 rpm were sub-cultured into fresh TSB and grown to OD600 of 1.0. Briefly, 100 μL of the bacteria were added to the wells and incubated at 37°C for 1 h. The plates were washed twice with 200 μL of PBS-T, fixed with 100 μL of 4% paraformaldehyde for 20 min at room temperature, washed twice with ddH_2_O and air-dried.

The adhered bacterial cells in the wells were stained with 150 μL of 0.1% Crystal violet and incubated at room temperature for 5 min. The plates were then washed twice with ddH_2_O and the crystal violet solubilized with 200 μL of 30% acetic acid at room temperature for 15 minutes. The absorbance was measured at OD595. The protein A null mutant *spa* was used as a negative control (Fig. S1).

### Bacterial binding to immobilized fibronectin and fibrinogen

Costar 96-well ELISA plates were coated with 10 μg of fibronectin (Sigma) for the fibronectin-binding assay and 10 μg of fibrinogen (Sigma) for the fibrinogen-binding assay in PBS overnight at 4°C. The wells were washed three times with PBS-T and air-dried for 1 hour. The plates were blocked with 1% BSA for 2 hours at 37°C.

Overnight cultures of *S. aureus* strains grown in TSB at 37°C and 200 rpm were sub-cultured into fresh TSB and grown to OD600 of 1.0. Briefly, an aliquot of bacteria was incubated at 37°C 10% FBS in TSB for 30 min. Following treatment, 100 μL of the bacteria suspension was added to the fibronectin-or fibrinogen-coated plate and incubated at 37°C for 1 h.

The plates were washed with 200 μL of PBS-T and fixed with 100 μL of 4% paraformaldehyde for 20 minutes at room temperature. The plates were washed twice with ddH_2_O after fixing and air-dried.

To visualize the attached bacterial cells, 150 μL of 0.1% Crystal violet was added to each well and incubated at room temperature for 5 minutes. The plates were washed twice with ddH_2_O and air-dried. To solubilize the crystal violet, 200 μL of 30% acetic acid was added to each well and incubated at room temperature for 15 minutes. The absorbance was measured at OD595. The fibronectin-binding protein AB mutant *fnbpAB* was used as a negative control (Fig. S1).

### Identification of two-component systems

The software HMMER (23) was used with the E-value cutoff of 0.01 to identify the two-component systems present in the whole-genome of the *S. aureus* USA300_FPR3757. The model for HMMER was created using sequences of two-component systems published in KEGG pathway database (https://www.genome.jp/brite/ko02022).

### Analysis of selection pressure in TCS genes

The analysis of site-specific selection pressure was analyzed using the method described by (24) with minor modifications. The *S. aureus* genomes were downloaded from the NCBI database. The Refseq, complete and annotated genomes were filtered based on deposited genomes between 2000 and 2023. The compiled database was composed of the CDS and the corresponding amino acid sequences in FASTA format. Orthologs of the TCS in *S. aureus* genomes were determined using the Orthofinder software (25) with the *S. aureus* USA300 TCSs as reference sequences. For each TCS HK and RR, duplicate sequences were removed using Seqkit program (26) and the orthologs were aligned by MUSCLE (27). The codon alignment was generated using pal2nal (28) and the aligned sequences were trimmed with trimAI (parameter:gappyout) (29). The treefile was generated from the aligned sequence using iqtree (parameter: st = DNA m = GTR+G4 nt = 1 fast) (30). To estimate mean posterior probabilities for selection pressure per site across each gene in the TCS was calculated by Fast, Unconstrained Bayesian AppRoximation for inferring selection (FUBAR)(31)

### Co-evolution analysis of TCS in *S. aureus* USA300

The analysis of coevolving residues between the DHp and the Rec domains was performed following computational methods described by (32) with minor modifications. The cognate pairs for all TCSs in the 1000 *S. aureus* genomes downloaded from the NCBI database were selected for the Orthofinder analysis using the cognate TCS pairs from the *S. aureus* USA300 as references. Duplicate TCS pairs were removed using the seqkit program(26). For the HKs, the transmembrane and sensor domains were removed and for the RRs, the output domains were removed. The remaining DHp and Rec domains were concatenated into a single sequence and aligned using MUSCLE (27). Columns in the alignment containing more than 10% gaps were eliminated from consideration. The mutual information analysis was performed using the online software MISTIC (33). The coevolving pairs were chosen based on an adjusted score of 3.5 or higher, ranking among the top 5 to 50 coevolving residues and those with the highest selection pressures. Although additional clusters of coevolving residues can be found at lower thresholds, this analysis focuses on the top 90% of coevolving residues within the cognate pairs.

### Protein modeling and phylogenetic analysis

The modeling of the *S. aureus* NarL HKs and RRs was generated with SWISS-MODEL (34). To determine the interacting residues between each NarL TCS cognate RR and HK, the models were superimposed onto the DesK-DesR cocrystalised structure (PDB: 5IUJ) (35) using PyMOL (36). Amino sequence alignments of the TCS RR and HK domains between different *Staphylococcus* strains and species, as well as *B. subtilis,* were performed with MUSCLE (27) and phylogenetic trees constructed with IQ-TREE (30).

### Statistical Analysis

Normality within the datasets was assessed using the Shapiro-Wilk test, where P-value <0.05 indicates the data do not follow a normal distribution. Subsequently, Levene’s test for equal variance was conducted to determine the appropriate test for comparing groups, either the Mann-Whitney U test for non-normally distributed data or Welch’s corrected T-test for normally distributed data with unequal variances.

## RESULTS

### Identification of two-component systems in *S. aureus*

To identify the TCSs in the *S. aureus* USA300, the genome was scanned using an HMMER model developed from the domains associated with TCSs from both gram-positive and negative bacteria. This revealed 16 TCSs belonging to NarL, OmpR, AraC, LytTR, and an unclassified family of TCS. Orthologs of the TCS were also identified in the *S. aureus* reference strain NCTC8325 and the MSSA476 (Table 1).

**Table 1.** Two-components systems identified in *S. aureus* USA300 and their orthologs in *S. aureus* reference strains NCTC8325 and MSSA476.

| Family | TCS | USA300_FPR3757<br>(SAUSA300_) | NCTC8325 ( SAOUHSC_) | MSSA476 |
| --- | --- | --- | --- | --- |
| AraC | HptSR | 0217/0218 | 00184/00185 | SAS0198/SAS0199 |
| NarL | VraSR | 1865/1866 | 02097/02098 | SAS1806/SAS1807 |
| NarL | NreBC | 2337/2338 | 02675/02676 | SAS2282/SAS2283 |
| NarL | DesKR | 1219/1220 | 01313/01314 | SAS1261/SAS1262 |
| NarL | Hypothetical | 1798/1799 | 01980/01981 | SAS1769/SAS1770 |
| OmpR | SrrAB | 1441/1442 | 01585/01586 | SAS1431/SAS1432 |
| OmpR | SaeSR | 0690/0691 | 00714/00715 | SAS0670/SAS0671 |
| OmpR | ArlSR | 1307/1308 | 01419/01420 | SAS1357/SAS1358 |
| OmpR | HssSR | 2308/2309 | 02643/02644 | SAS2252/SAS2253 |
| OmpR | GraSR | 0645/0646 | 00665/00666 | SAS0624/SAS0625 |
| OmpR | BraSR(NsaSR) | 2558/2559 | 02955/02956 | SAS2510/SAS2511 |
| OmpR | VicKR(WalKR) | 0020/0021 | 00020/00021 | SAS0018/SAS0019 |
| OmpR | PhoPR | 1638/1639 | 01799/01800 | SAS1620/SAS1621 |
| OmpR | KdpDE | 2035/2036 | 02314/02315 | SAS1983/SAS1984 |
| LytTR | AgrAC | 1991/1992 | 02264/02265 | SAS1943/SAS1944 |
| LytTR | LytST | 0254/02545 | 00230/00231 | SAS0237/SAS0238 |
| Unclassified | YycHI | 0022/0023 | 00022/00023 | SAS0020/SAS0021 |

Putative domain search using SMART (37) showed the presence of domains such as the Transmembrane (TMR), HisKA (dimerization and phosphoacceptor) and the HATPase (Histidine kinase-like ATPase) domains in most of the HKs. Some of the HKs contained additional domains such as the HAMP, PAS, GAF, and DUF4118 domains commonly found between the transmembrane and the HisKA domains in HKs (Fig. 1A). The unclassified HK YycI possessed a transmembrane domain and a YycI domain whereas the AgrC possessed six TMR and a HATPase domain. The NreB and the SAUSA300_1799 lacked the TMR whereas the 4 predicted TMR in the KdpD overlap with DUF domain. The response regulators (RRs) contained the common Rec (phosphoacceptor) domain in all the RRs with either of the effector binding domains such as the Trans Reg, LytTR, HTH-LUXR, and AraC in different RRs. These domains were absent in YycH (Fig. 1B).

**FIG 1.**
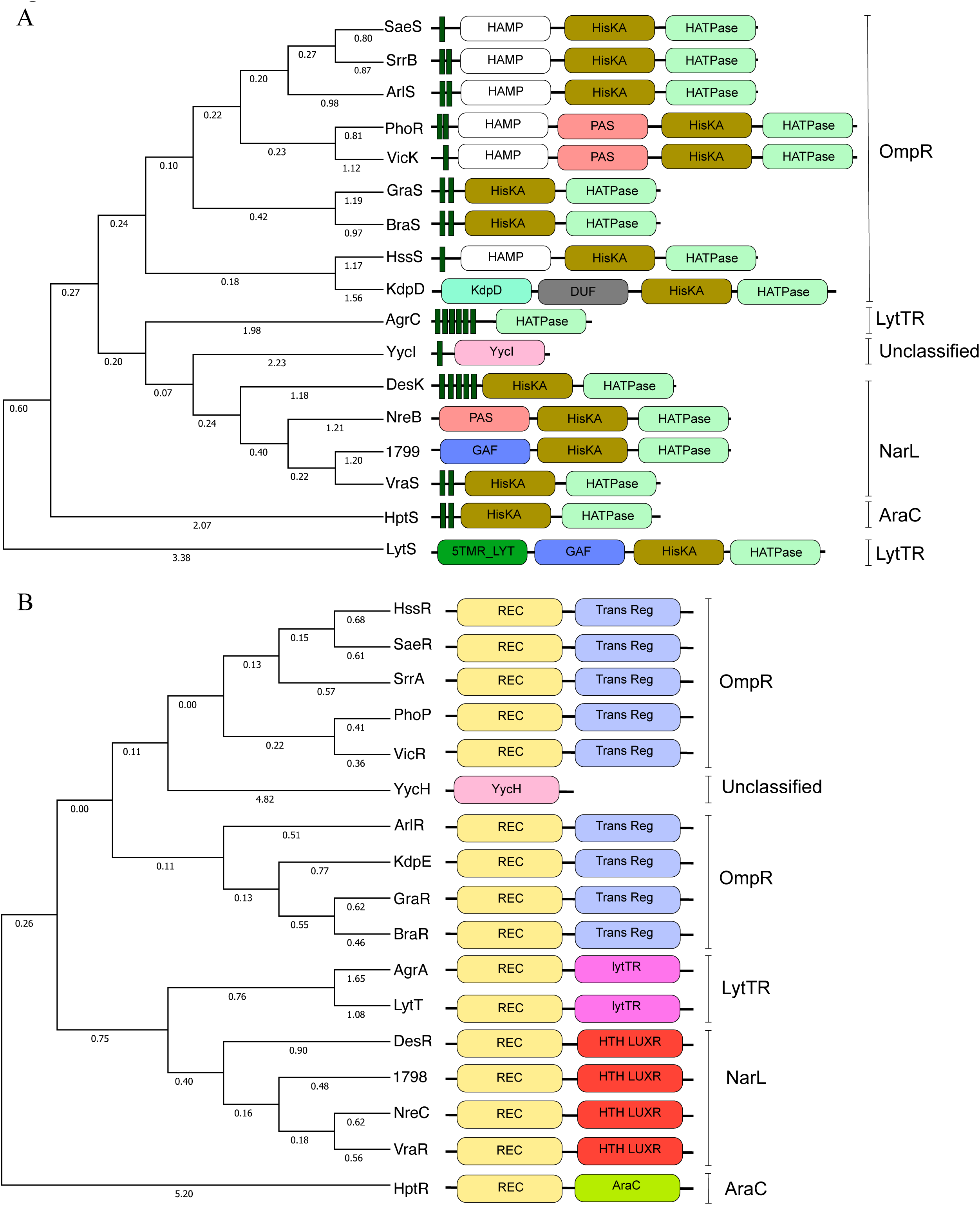
Domain architecture and phylogenetic analysis of the *S. aureus* USA300 TCS: **A**) Histidine Kinases (HKs) and **B**) Response Regulators (RRs). Phylogenetic analysis was performed using the amino acid sequences of the components from the USA300-FPR3757 genome, employing maximum likelihood testing and a bootstrap value of 1000. Domains corresponding to the TCS components are indicated by labelled, colored rectangular blocks, while vertically labelled lines show families.The Transmembrane (TMR) domains of the HKs are shown as narrow green rectangular blocks.

Phylogenetic analysis based on the amino acid sequences of the TCSs, demonstrated high relatedness between the HKs and RRs within their respective families. This clustering was more evident in the RRs, as they share more conserved domain architecture. Interestingly, even though some HKs possessed additional domains, they still exhibited close relatedness within their families, except for the LytTR and unclassified HKs (Fig. 1). This observation suggests that, despite the presence of extra domains, the core structural features of these TCSs are conserved across their respective families.

### Impact of TCSs on virulence of *S. aureus*

The common domain architecture and relatedness between individual HKs and RRs within TCSs can influence their regulation of virulence determinants and lead to possible cross-regulation of biological pathways. To investigate how the conservation and clustering of HKs and RRs influence their control of virulence determinants, the transposon mutants of the TCS components were examined for their effect on biofilm formation, hemolytic activity, protease activity, protein A production, nuclease production, fibronectin and fibrinogen binding, and the ability to infect, adhere and kill human cell lines.

Mutants of TCSs such as YycIH, AgrAC, LytTS, NreBC, BraRS, ArlRS, PhoPR, HptRS, and VraRS were significantly less cytotoxic for HaCaT cell lines than the WT (Fig. 2A). In contrast, the GraSR, HssR and KdpE mutants displayed increased cytotoxicity, while the remaining TCS mutants showed either marginal reductions or no significant differences in cytotoxicity when one of the components was inactivated. In some of the TCS, mutations in either the HK or the RR resulted in varying effects on their cytotoxicity. For instance, in SaeRS, the absence of the HK significantly reduced cytotoxicity, while gene inactivation of its cognate HK had a similar effect as the WT. These observations may suggest the possibility of crosstalk activities among different HK or that the RR SaeR can also respond to other regulatory systems (Fig. 2A).

**FIG 2.**
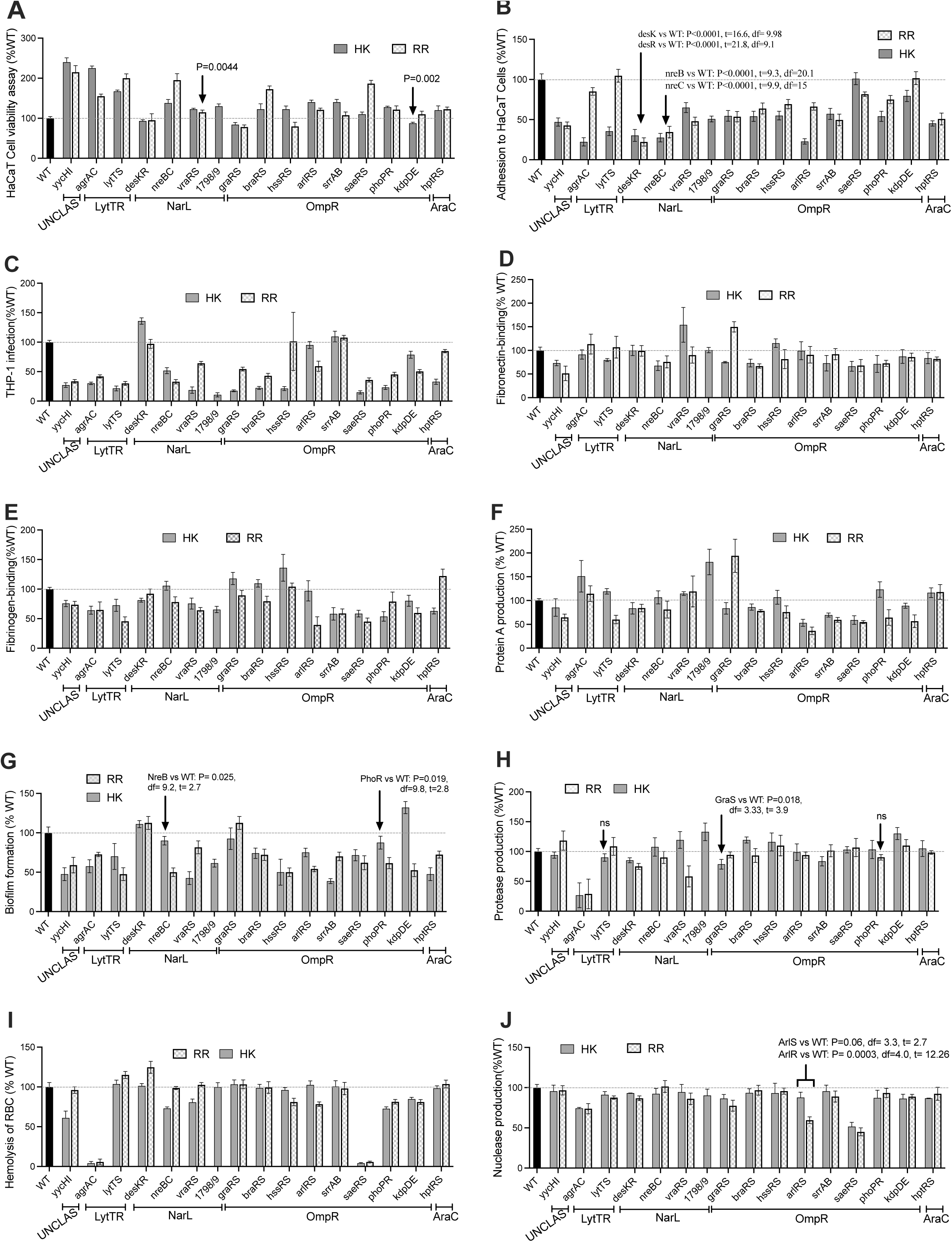
Virulence phenotype of TCS component mutants. Virulence profiles of the HKs and RRs mutants grouped based on the family. **A**) HaCaT cell viability assay, **B**) adhesion of bacteria to HaCaT cells, **C**) bacterial infection of THP-1 cells, **D**) Fibronectin, **E**) Fibrinogen binding, **F**) protein A production, **G**) biofilm formation, **H**) protease production **I**) hemolysis of red blood cells and **J**) nuclease production of the *S. aureus* TCS component transposon mutants. The cognate pair of HKs and RRs are grouped based on their respective family. The production of the virulence phenotypes is expressed as a percentage of the wildtype (WT) normalised to 100%. All data represent mean± SD of n=4 independent experiments.P-value <0.05 is considered significant.

Moreover, adhesion of *S. aureus* to HaCaT cells was generally diminished among the TCS mutants, except for HK SaeS and the RRs LytT and KdpE (Fig. 2B). The DesKR, NreBC showed the most significant reduction in adhesion to HaCaT cells. Additionally, compared to the WT, the TCS mutants mostly exhibited reduced infection in THP-1 cell lines except for the DesKR, SrrAB, HssR, and ArlS (Fig. 2C).

*S. aureus* cell wall anchored factors facilitate bacterial attachment to host tissue, promoting colonization and evasion of host immune factors (2, 3). While previous reports have indicated that SaeRS regulates fibronectin and fibrinogen binding (38, 39), we observed reduced binding to immobilized fibronectin in the mutants of TCSs YycHI, NreBC, BraSR, and SaeRS (Fig. 2D). In contrast, mutants of AgrAC, DesKR, VraSR, HssSR, and ArlSR showed no difference or increased binding to fibronectin compared to the WT. Distinct variations in binding to fibronectin were observed among some mutant RRs and HKs, including LytS, HptR, and PhoR (PhoR vs WT: P=0.09, df=2.6, t=2.14) when compared to their corresponding cognate pairs (Fig. 2D). A particularly significant deviation was noted within the GraSR system’s HK-RR pair. The GraR mutant (GraR vs WT: P=0.004, df= 3.5, t=-6.5) demonstrated increased binding to fibronectin compared to the WT, whereas the GraS mutant (GraS vs WT: P=0.01, df=2.2, t=6.12) revealed a diminished binding compared to the WT.

From our assay, only YycHI and SaeRS influenced both adhesions to immobilized fibrinogen and fibronectin (Fig. 2D and E). Additionally, we observed decreased binding in the SrrAB, AgrAC, LytTS, VraRS, and KdpDE compared to the WT (Fig. 2E). The AgrAC, typically a negative regulator of fibronectin-binding (40)(Fig. 2D), exhibited a positive regulatory effect on fibrinogen-binding. Interestingly, both HK-RR mutants of HssRS showed a marginal increase (HssS vs WT: P=0.01, df=5.1, t=-3.8) or WT-level (HssR vs WT: P=0.19, df=7.48, t=-1.2) binding to fibrinogen, while individual response regulators (RRs) HtpR and DesR, and HKs BraS, NreB, and ArlS exhibited increased or similar binding profiles to fibrinogen compared to the WT (Fig. 2E). Although HssRS is known for its role in sensing heme concentration, there is no direct experimental evidence of its influence on fibrinogen and fibronectin binding. HssRS controls *htr*A expression, which counteracts heme toxicity. Mutations in *hssR* and *htrA* lead to *S. aureus* hypervirulence via heme buildup, while *htrA* mutants upregulate fibronectin and fibrinogen-binding proteins (41–43), which could explain the increased binding of HssRS mutants to fibrinogen and fibronectin (Fig. 2D and E).

In addition to the SaeRS and ArlRS, which positively regulates bacterial surface-associated Protein A (44), mutations of SrrAB showed a significant decrease in binding to immobilized anti-Protein A antibodies compared to the WT (Fig. 2F). This is possibly due to the SrrAB negative regulation of the *agr* locus (45). In agreement, the mutation of the HK-RR of AgrAC showed increased binding to the immobilized anti-protein A antibodies (44)(Fig 2F). The mutants of the HK-RR pairs of KdpDE (KdpD vs WT: P=0.05, df= 3.7, t= 2.8) and the BraSR (BraS vs WT: P=0.04, df= 3.32, t= 3.12) showed a marginal difference in binding to anti-protein A antibodies. VraRS, HptRS, and the individual RR GraR and HKs LytS, SAUSA300_1799, HssS, and PhoR displayed increased binding to anti-protein A antibodies compared to the WT (Fig. 2F).

Biofilm formation in *S. aureus* is a complex and dynamic process regulated by a network of interconnected TCSs. Mutation of the TCSs, except for DesKR, GraSR, the HK KdpD were found to reduce biofilm formation (Fig. 2G). TCSs like ArlRS, AgrAC, SrrAB, SaeRS, and LytTS play critical roles in modulating the expression of surface proteins, extracellular matrix components, and quorum-sensing molecules, which contribute to biofilm initiation, maturation, and dispersal (46).

While the influence of GraS on protease production is most notable under acidic pH conditions (47), we found a 15-20% reduction in protease activity for the *graS* mutant compared to the WT in cation-adjusted MH at pH7. Interestingly, there was no significant difference in protease production between the *graR* mutant and the WT (Fig. 2H). Although ArlRS and SaeRS have been reported to control various genes in protease loci (48), our analysis showed no impact on protease production for their corresponding mutants. In contrast, TCS AgrAC mutants demonstrated the most substantial effect on protease production. Furthermore, mutations in both DesKR components led to a 15-20% decrease in protease production compared to the WT level. Additionally, while the VraR mutant exhibited a significant reduction in protease production, its sensor kinase VraS showed no discernible difference from the WT (Fig. 2H).

Hemolysis was most notably reduced in the AgrAC and SaeRS mutants. Additionally, the PhoPR and KdpDE mutants displayed decreased hemolysis compared to the WT (Fig. 2I). Furthermore, mutants of the HKs YycI, NreB, VraS, and the RRs HssR, ArlR showed reduced hemolysis compared to their cognate pairs, which were similar to WT levels (Fig. 2I). Both SaeRS and AgrAC TCSs are known to influence hemolysis in *S. aureus* (39). Although ArlSR has been shown to indirectly affect hemolysis by regulating SaeRS via ScrA (49), our study only observed reduced hemolysis in the ArlR mutant (ArlR vs WT: P<0.0001, df=7.6, t=8.6) (Fig. 2I).

Nucleases play a critical role in evading the host immune system by degrading host DNA, as well as in nutrient acquisition and biofilm maintenance (50). Among the TCS mutants, SaeRS exhibited the most significant decrease in nuclease production, followed by ArlR, AgrC, and, to a lesser extent, the GraRS system (Fig. 2J). This is in agreement with previous studies that demonstrated the influence of SaeRS and ArlRS on nuclease production (51, 52).

Considering the regulatory impacts of the TCS components on virulence, we explored TCS pairs that exhibit similar cross-regulation of virulence profiles. Principal component analysis (PCA) of the virulence profiles in the HK and RR mutants identified five clusters that displayed limited similarity in their family classifications. Cognate pairs that showed similarities in virulence regulation included the SrrAB, SaeRS, YycHI, BraSR, PhoPR, NreBC, AgrAC, and DesKR (Fig. S1A and B). Among these gene sets, the SaeRS and AgrAC have global regulons and influence a subset of genes controlled by other TCSs in *S. aureus* (53). Furthermore, the AgrAC formed a separate cluster, suggesting its ability to influence the production of one or more virulence determinants independently or significantly. Despite belonging to the same family as LytTR (Fig. 1), the AgrAC did not cluster with LytT or LytS (Fig. S2). HKs, including KdpD, ArlS, and DesK, and the RRs HptR, HssR, and DesR, demonstrated similar virulence regulatory profiles as the WT. These results suggest that possible cross-regulation between the TCS in regulating biological pathways is crucial for the bacterium’s infection potential. Also, the observed variations in virulence regulatory profiles among TCS components and the clustering noted from the PCA show significant crosstalk among different TCSs. Such interactions amplify or mitigate the bacterium’s virulence, shaping the overall pathogenic potential of *S. aureus*.

### Evolution and selection of TCS in *S. aureus* strains

The role of TCS in virulence regulation and their conservation in bacterial genomes make them potential targets for antimicrobial agents. Investigating selection pressure across TCS gene sites provides insights into the evolutionary constraints and functional diversity of the TCS. To further explore the selection pressure, evolutionary stability, and conservation of TCSs in *S. aureus* strains, we conducted an evolutionary test of selection pressure using phylogenies of TCSs derived from 1000 complete and annotated *S. aureus* genomes. For this analysis, we used FUBAR, which applies the Bayesian approach to infer nonsynonymous (dN) and synonymous (dS) substitution rates on a per-site basis for a given coding alignment and corresponding phylogeny (31). Orthologs of each TCSs RR and HK from the 1000 *S. aureus* genomes (from 2000 to 2022 available at NCBI) were used for codon alignment and phylogeny. More than 86% of the analyzed genomes contained all TCSs present in USA300_FPR3757, indicating high evolutionary conservation (Fig. S3).

Overall, we noted more pronounced negative (purifying) selection pressures on the HKs in comparison to the RRs (Table. S2). On the other hand, 7 HKs and 9 RRs were shown to undergo diversifying (positive) selection with 1 to 3 sites under positive selection pressure (Fig. 3 and Table S2).

**FIG 3.**
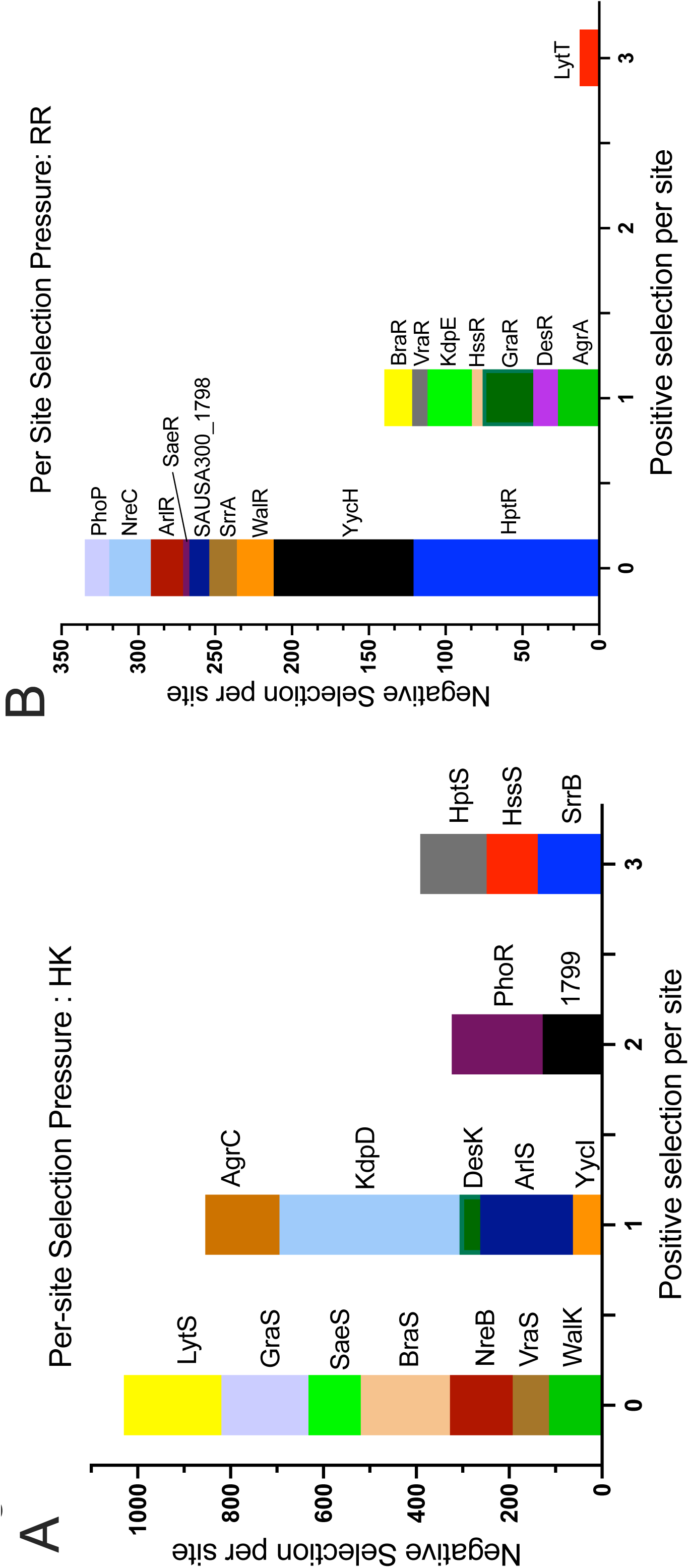
The selection pressure per-site in the HKs and RRs of the TCS in *S. aureus*. The number of sites under negative selection pressure (y-axis) and positive selection pressure (x-axis) in the **A**) HKs and **B**) RRs of the TCS from the *S. aureus* genomes.

Furthermore, no positive (diversifying) selection was identified for residues in HKs VicK, VraS, NreB, GraS, BraS, and LytS. One site under positive selection pressure was identified in the YycI, ArlS, DesK, SaeS, KdpD, and AgrC, while SAUSA300_1799, PhoR, SrrB, HssS, and HptS had 2 or 3 sites undergoing positive selection pressure (Fig. 3A and Table. S2).

Fewer sites under positive and negative selection pressures were observed in the RRs compared to their cognate HKs. Most RRs, including PhoP, HptR, WalR, YycH, ArlR, SrrA, SAUSA300_1798, SaeR, and NreC, had no residues under positive selection pressure (Fig 4B and Table S3B). In contrast, RRs GraR, HssR, KdpE, VraR, BraR, AgrA, and DesR had one site with inferred positive selection, while LytT had two sites with positive selection pressure (Fig. 3B and Table. S2). In both the HKs and RRs, no sites under positive selection were observed in the TCSs WalKR, NreBC, and SaeRS. This finding aligns with expectations for WalKR, an essential TCS (54). Conversely, while NreBC and SaeRS are not deemed essential TCSs for bacterial viability, their cognate pairs exhibit high specificity in regulating virulence determinants (Fig 3 and Table S2).

**FIG 4.**
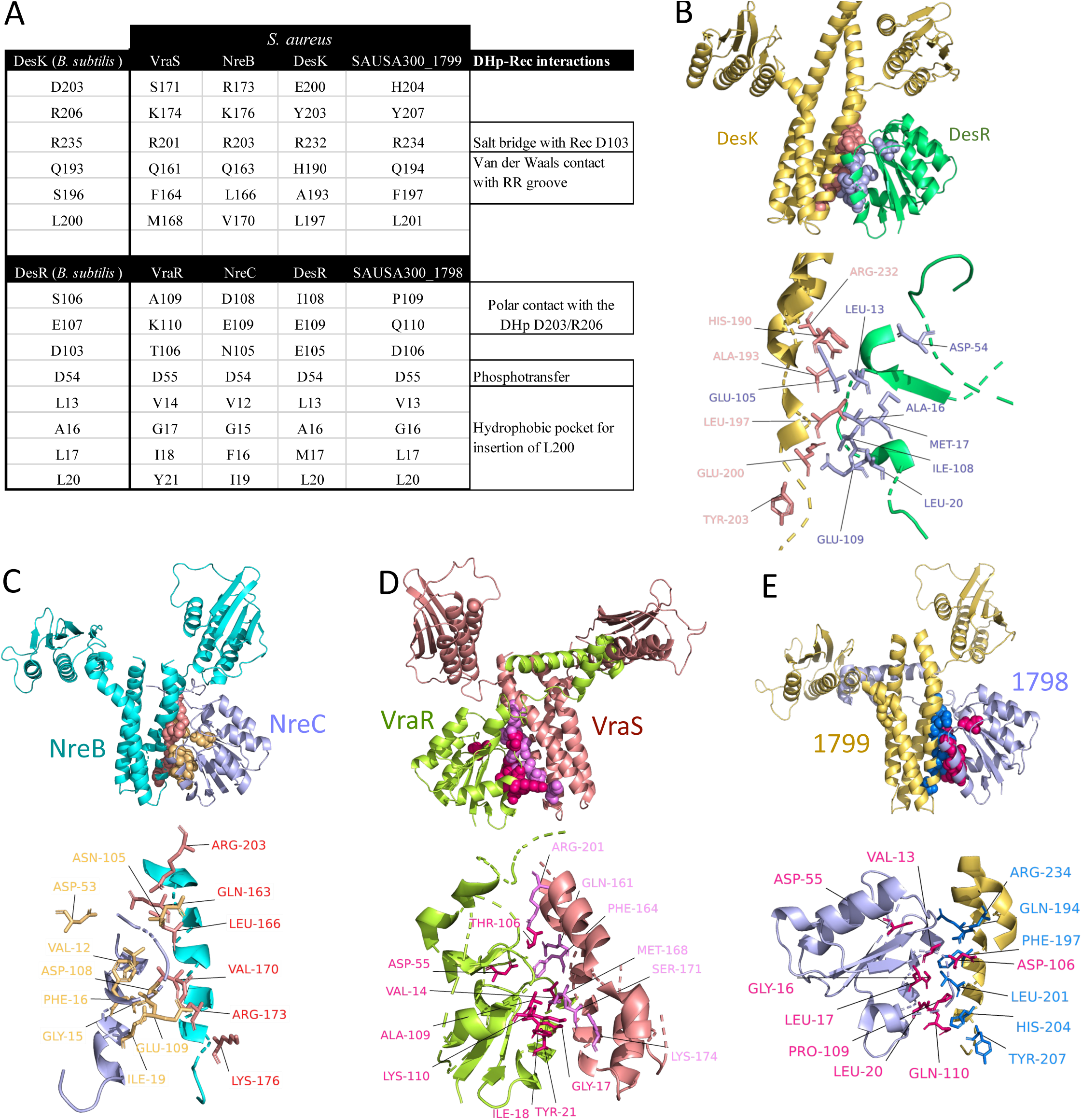
**A**) Interacting residues from *B. subtilis* DesK and DesR, as well as *S. aureus* HKs and RRs from the NarL family, are displayed. The selected residues from *S. aureus* are based on their alignment with the interacting residues of DesK (DHp) and DesR (Rec) domains. Each residue is shown with its amino acid and position within the specific protein. **B**) Superimposing the RR and HK domains from the modelled *S. aureus* DesK and DesR onto the cocrystal structure of the *B. subtilis* DesKR demonstrated high similarities in alignment (1.993Å RMSD aligning 2296 atoms of the HK and 0.351Å RMSD aligning 626 atoms of the RR). **C)**. The NreB-NreC model using the *B. subtilis* DesKR revealed very similar structures, with 2.114Å RMSD aligning 1959 atoms of the HK and 0.402Å RMSD aligning 601 atoms of the RR. **D**) Superimposition of the modelled VraRS with the *B. subtilis* DesK-DesR complex showed high similarity in alignment, with 2.487Å RMSD aligning 2006 atoms of the HK and 0.322Å RMSD aligning 577 atoms of the RR. **E)** Superimposition of the modelled SAUSA300_1798/9 with the *B. subtilis* DesK-DesR cocrystal structure showed high similarity in alignment, with 2.114Å RMSD aligning 1959 atoms of the HK and 0.402Å RMSD aligning 601 atoms of the RR.

### Coevolution of specificity determinants in NarL TCS

Given the selection pressures across the sites of HKs and RRs and the fact that certain cognate RR and HK mutants can independently maintain virulence phenotypes, we aimed to further elucidate the evolutionary relationship between the interacting residues of cognate TCS pairs. Given the importance of HK-DHp and RR-Rec domain residue interactions in TCS specificity (32), we investigated the evolutionary link between these interacting residues in TCS components, exploring the potential for TCS crosstalk.

We employed the NarL TCS DesKR cocrystal structure from *Bacillus subtilis*_168 (35) as a modelling proxy to illustrate the variations in specificity-determining residues between the DHp and Rec of the NarL TCS in *S. aureus* consisting of DesKR, VraRS, NreBC, and SAUSA300_1798/1799. Among these members, NreBC, VraS, and SAUSA300_1798 exhibit no sites under diversifying selection (Fig 3 and Table S2 which suggests strong evolutionary stability and indicates functional constraint. Additionally, cognate pairs of the DesKR and the NreBC showed similarities in the control of virulence determinants, which signifies the conservation of signal transduction within the TCSs. Contrastingly, the HK VraS and its associated RR VraR displayed divergent virulence regulatory profiles, suggesting potential crosstalk with other systems (Fig 2 and S2). While VraS and SAUSA300_1799 exhibit similar virulence profiles, the lack of a transposon mutant for RR SAUSA300_1799 hinders a comprehensive comparison for potential cross-regulation in virulence between the two system. Through amino acid sequence alignment between *B. subtilis* DesKR and *S. aureus* NarL TCSs, we mapped residues in the DHp (6 residues) and Rec (8 residues) domains that mediate interactions. The interacting residues varied between the various *S. aureus* NarL TCSs and the DesKR from *B. subtilis* (35)(Fig.4A, S4, and S5). Superimposition of the *S. aureus* NarL TCSs demonstrated a similar localization of interacting residues within the DHp and Rec domains, consistent with the arrangement observed in the *B. subtilis* DesKR cocrystal structure (Fig. 4B-E and S6). We did not analyze polar interactions between the residues, as these models are only derived from alignment with predetermined structures and may not provide accurate distances.

Investigation of the conservation of interacting residues among the NarL TCSs in other *Staphylococcus* species (*S. aureus* Newman, *S. aureus* USA300, *S. aureus* MSSA476, *S. epidermidis*, *S. aureus* NCTC8325, and *S. haemolyticus* ATCC14990), and *B. subtilis* revealed that within the same bacteria, interacting residues of DHp and Rec domains vary among different NarL TCSs (Fig 5 and 6), which implies evolution of distinct specificities for cognate partners. Also interacting residues of orthologous TCSs were observed to be highly conserved, even across different strains and species of *Staphylococcus* (Fig. 5 and 6), highlighting the evolution of functional conservation.

**FIG 5.**
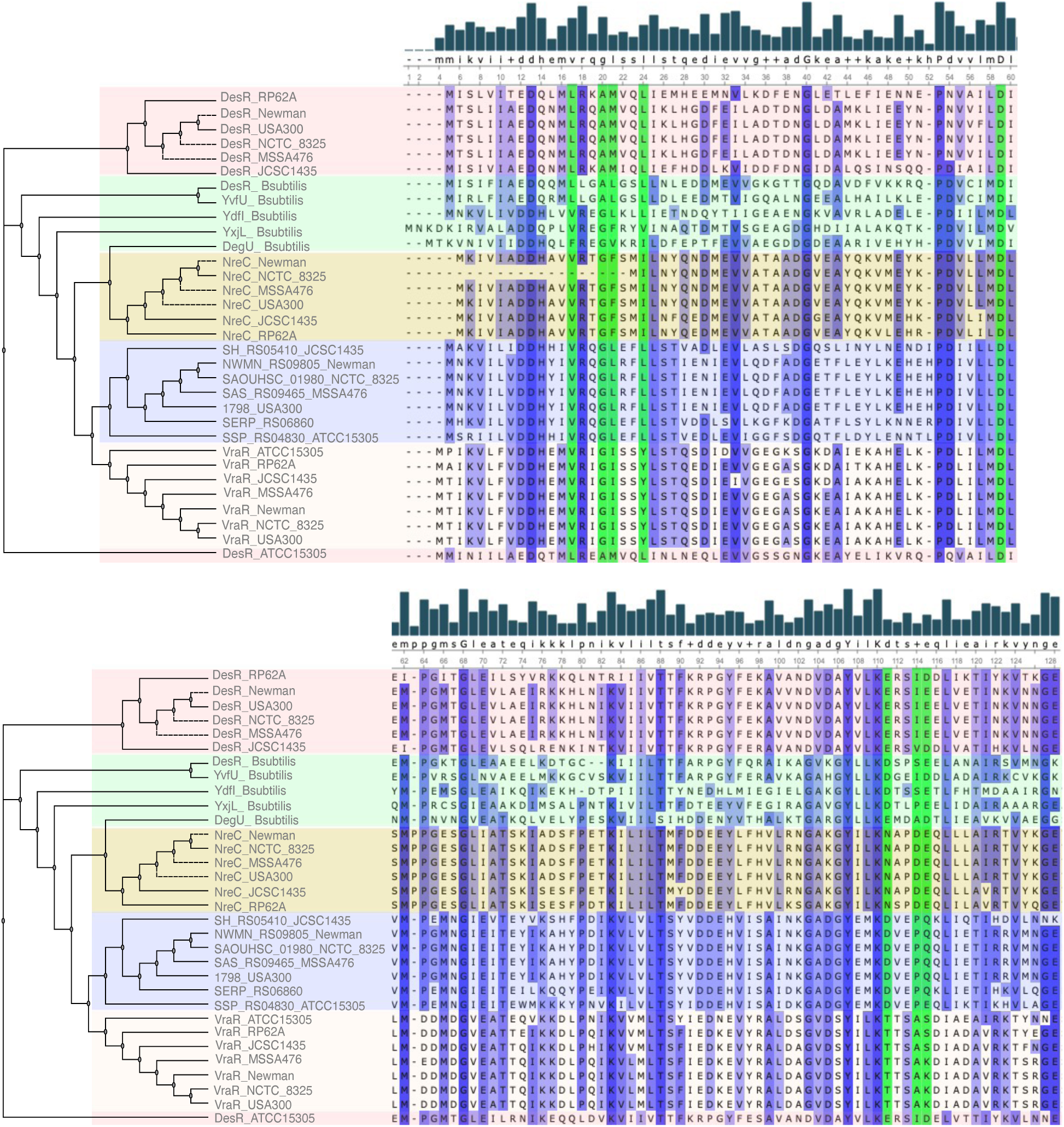
The phylogenetic analysis and sequence alignment illustrate the relationships between the NarL TCS RR Rec domains from *Staphylococcus* spp. and *B. subtilis*. Gaps introduced in the sequence alignment serve to maximize alignment and are represented as dashes. The green vertical shading indicates the aligned residues that interact with the DHP domain. The blue vertical shading, with increasing intensity, signifies conserved regions. The degree of consensus sequence is depicted by the bar chart, with uppercase residues representing highly conserved areas and lowercase residues for less conserved regions. The horizontal colored shading demonstrates the clustering based on the percentage identity matrix of the aligned amino acid sequences of the full-length RR from each TCS (Fig. S4B). Groups under the same horizontal colored shading belong to the same cluster. Phylogenetic analysis was performed with MUSCLE with Bootstrap value =1000.

**FIG 6.**
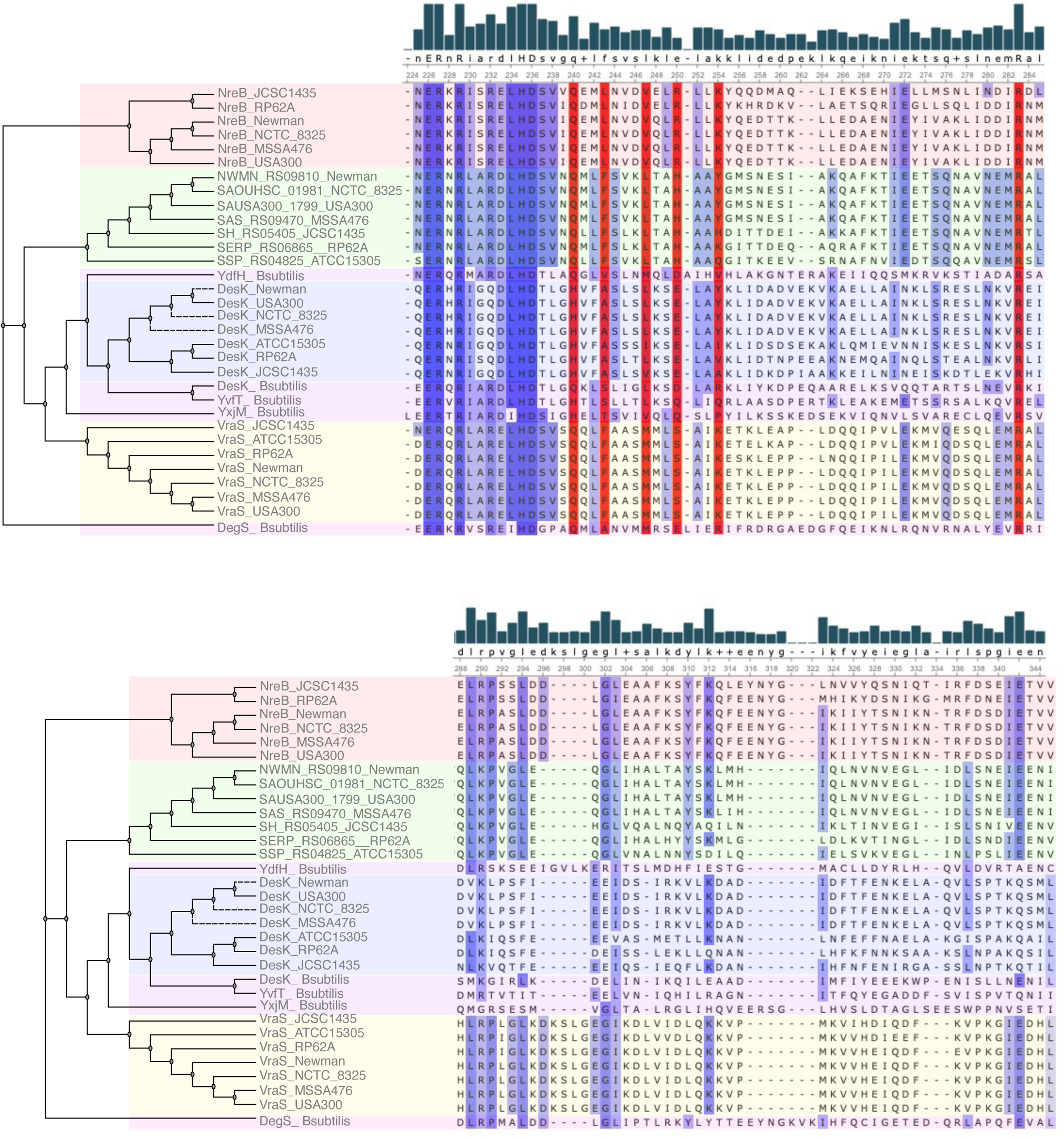
The phylogenetic analysis and sequence alignment illustrate the relationships between the NarL TCS DHp domains from *Staphylococcus* spp. and *B. subtilis*. Gaps introduced in the sequence alignment serve to maximize alignment and are represented as dashes. The red vertical shading indicates the aligned residues that interact with the rec domain. The blue vertical shading, with increasing intensity, signifies conserved regions. The degree of consensus sequence is depicted by the bar chart, with uppercase residues representing highly conserved areas and lowercase residues for less conserved regions. The horizontal colored shading demonstrates the clustering based on the percentage identity matrix of the aligned amino acid sequences of the full-length HK from each TCS (Fig. S4A). Groups under the same horizontal colored shading belong to the same cluster. Phylogenetic analysis was performed with MUSCLE with Bootstrap value =1000.

The coevolution of interacting residues in cognate TCS pairs is fundamental to ensure specificity and reduces unwanted crosstalk with other TCS components (35). To gain further insight into the evolution of TCSs in maintaining signal fidelity between cognate HK-RR pairs, we analyzed the coevolution of residues between the Rec and DHp domains of cognate NarL TCSs. This was done by Mutual information analysis of the amino acid alignments in cognate HKs and RRs of *S. aureus* (Fig S7). In the *S. aureus* DesKR’s DHp and Rec domains, we did not identify a significant correlation between residues indicative of coevolution. This observation aligns with our expectations, given the low selection pressure on the DesKR HK and RR (Fig. 3 and Table S2). Notably, among the interacting residues, a negative selection pressure was observed at the DHp residue R232_DesK_. This residue, conserved across NarL HK DHp domains (Fig. 6), forms a salt bridge with the *B. subtilis* D103_DesR_ (35).

Analysis of coevolutionary patterns in the DHp and Rec domains of NreBC highlighted a correlation between residues K176_(DHp)_ and N105_(Rec)_, both of which play a role in the interaction between these domains. Referring to the *B. subtilis* DesK-DesR cocrystal structure, K176_(DHp)_ establishes polar contacts with D108_(Rec)_ and E109_(Rec)_. In contrast, N105_(Rec)_ forms a salt bridge with R203_(DHp)_ (35). Additionally, D108_(Rec)_ and E109_(Rec)_ exhibit a strong correlation with T181_(DHp)_ and L172_(DHp)_, respectively. Furthermore, the residue Q163_(DHp)_, which creates van der Waals contacts within the RR groove, is highly correlated with A98_(Rec)_ (Fig. 7A). Among the coevolving residues identified, those from the DHp domain, including E149, K151, K183, L185, L214, Q163, R210, and S154, as well as those from the Rec domain, namely G97, I78, and P74, were deduced to be under negative selection pressure (Fig. 7A). Consistent with expectations, none of the residues that interact between the DHp and Rec domains exhibited positive selection pressure (Fig. 7A).

**FIG 7.**
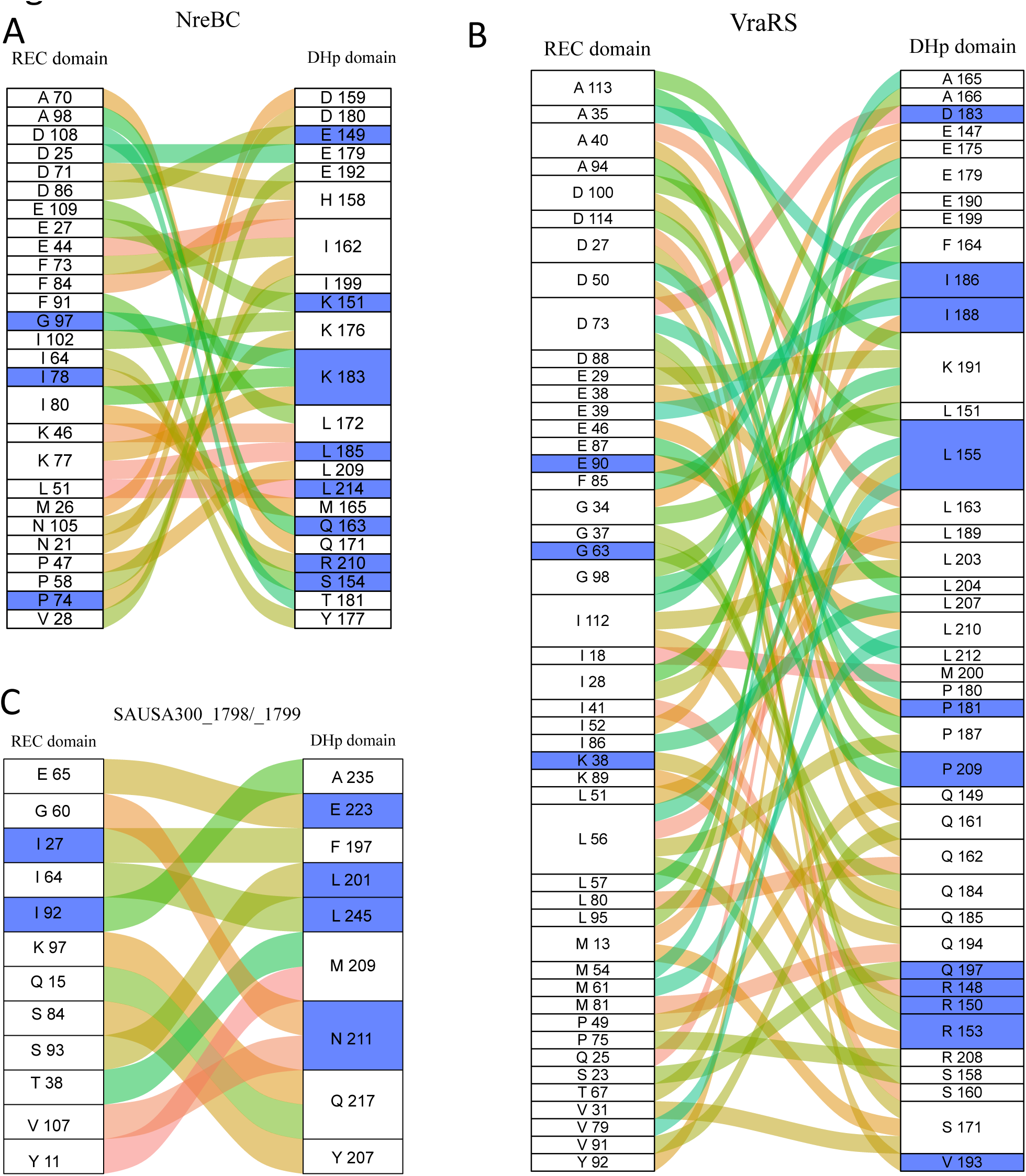
The coevolving residues of the Rec and DHp domain of the *S. aureus* **A**) NreBC, **B**) VraSR and **C**) SAUSA300_1798/1799. Curved connecting lines Curved lines show correlations between Rec and DHp residues. Blue shaded boxes represent residues that show evidence of posterior probability for negative selection. A threshold of 0.9 is set for inferring positive or negative selection.

In the VraRS TCS, the interacting residues Q161_(DHp)_, F164_(DHp)_, S171_(DHp)_ and the I18_(Rec)_ were noted to coevolve with other residues in the cognate pair domains (Fig. 7B). Despite these coevolutionary tendencies, there was no evident coevolution between directly interacting residue pairs, and no selection pressures (neither positive nor negative) were identified for these residues or their coevolving counterparts (Fig. 7B).

In the SAUSA300_1798/1799 TCS, the residue F197 _(DHp)_, which forms van der Waals interactions with the RR groove, notably coevolves with I27_(Rec)_ that is under negative selection pressure. Intriguingly, the L201_(DHp)_ residue, aligning with the hydrophobic groove of the Rec domain, strongly correlates with S93_(Rec)_. Further emphasizing its evolutionary significance, L201_(DHp)_ exhibits evidence of negative selection pressure (Fig 7C).

In general, the presence of negative selection pressures on various residues emphasizes their pivotal roles in system functionality and suggests evolutionary constraints to conserve these crucial interactions. Interestingly, while some TCSs showed discernible coevolution patterns, others did not, reflecting diverse evolutionary strategies or the influence of different environmental pressures.

## DISCUSSION

In this study focusing on *S. aureus USA300*, a predominant MRSA lineage, we unravel the complex interplay between the TCSs and uncover shared and unique virulence regulatory pathways and the genetic variations that mediate their signal specificity. Here, a comprehensive investigation of the virulence regulation of the TCSs was conducted using a transposon mutant library of the USA300 JE2 strain carrying inactivated HKs and RRs. We observed that some TCSs from the same family regulated different functions (Fig. 2 and S2), which was expected due to their divergent functional evolution for sensing and responding to specific stimuli. Functional differences among TCS can be linked to the diversification of sensor and phosphotransfer domains in the HKs, where closely related TCS with similar domain architecture may show variations in specificity to cognate and non-cognate RRs (8, 18). Also, the modular architecture of the TCS, which includes signal sensing, phosphotransfer, and response, can contribute to variations in the impact of different TCSs on virulence determinants (8). Moreover, the integration of TCS within larger regulatory networks enables parallel or hierarchical regulation of virulence mechanisms or modulation by other active TCSs. For example, multiple interconnected TCSs such as ArlS, AgrAC, SrrAB, SaeRS, and LytT regulate biofilm formation (Fig 2G).

While profiling the virulence of the TCS mutants, we did not use complements of the transposon mutants due to inherent technical and genetic constraints with the maintenance of modified vectors in the strains without antibiotic induction, which will alter the conditions for the virulence profiling creating discrepancies in the analysis. Instead, we compared the impact of the mutants against the WT and correlated our findings with existing published studies to draw meaningful conclusions on their virulence attributes. This study did not factor in the potential for non-specific effects arising from secondary mutations or polar effects in the transposon mutants. To mitigate this, we selectively focused on transposon mutants with insertions impacting the genes and restrained our selection to those with insertions within the first 60% of the gene.

Our study found that several TCSs exhibit similar regulatory profiles on virulence factor production despite belonging to different TCS families. This suggests divergent regulatory pathways have evolved despite similarities in HK and RR domains. Notably, global regulators AgrAC, SrrAB, and SaeRS displayed a high degree of correlation in their virulence profiles. The ArlRS and SaeRS TCSs indirectly influence each other’s activity through their connection with the Agr quorum sensing system, which is strain-dependent and affected by environmental conditions and the growth phase (4, 55, 56). The virulence profiles reveal an overlapping regulon between SaeRS and AgrAC, but SaeRS shares more common virulence profiles with SrrAB (Fig. S2). Both SaeRS and SrrAB belong to the OmpR family (Fig. 1).

We also identified cognate TCS pairs with similar virulence gene profiles, including DesKR, BraSR, PhoPR, NreBC, and YycHI, which can imply the evolution of specialized functions for these TCSs. Fascinatingly, the mutations of the HKs in the BraSR and GraSR displayed similarities in virulence profiles, which matches their similar domain architecture and sequence homology. However, their RR mutants exhibit distinct virulence profiles, suggesting functional divergence in RRs regulating downstream pathways.

Potential instances of independent functionality (e.g., AgrAC), crosstalk (e.g., ArlS-DesR and SrrB-VraR), and evolutionary divergence toward global regulon and specialized functionality (e.g., AgrAC and SaeRS) were observed among some of the TCSs. It is important to note that our assays used primary growth media and may lack the stimuli to activate TCS, resulting in less virulence factor production variation than previously published findings. For example, SrrAB exhibits both positive and negative regulation of protein A production under aerobic (Fig. 2F) and microaerobic conditions, (57) respectively.

The TCS interconnections within broader regulatory networks enable parallel or hierarchical regulation of virulence mechanisms or modulation by activity of other TCSs (58–60). For instance, multiple interconnected TCSs like ArlS, AgrAC, SrrAB, SaeRS, and LytTS regulate biofilm formation. This cross-talk among TCSs leads to complex and dynamic virulence factor regulation, impacting bacterial adaptation and pathogenicity. Crosstalk between TCS from the same family is expected due to common ancestry (5); however, it can also occur between different families (12). This depends on the evolution of specificity determinants mediating HK-DHp and RR-rec domain interactions for phosphotransfer. Given the selection pressure on TCS, continuous evolution could rewire signaling pathways or insulate TCS pathways through accumulated mutations in specificity determinants.

This study used modelled structures of the NarL family TCS and the cocrystalized DesK-DesR complex from *B. subtilis* (35) to investigate the interacting residues between cognate TCS pairs. Sequence alignment revealed variations in these residues across different NarL TCSs from *S. aureus* strains and species. This highlighted the evolution of specificity determinants between cognate TCS pairs (8, 18), essential for reducing unwanted crosstalk and evolving new functions. We concurrently observed the conservation of specificity determinants within the same TCS across multiple strains. We also observed variations among related TCS within the same family from other bacterial species. While mutations in residues near interaction interfaces could influence specificity, we focused on the residues at the interaction interface between the HKs and RRs of *S. aureus* NarL TCS, referencing the DesK-DesR TCS from *B. subtilis*. Notably, we found species-specific variations in the interacting residues within the TCS belonging to the same family. This means that single amino acid changes in HK residues involved in specificity could lead to altered interactions with the RR as previously experimentally determined (8, 61, 62). Crucial similarities in biochemical properties, such as polarity and hydrophobicity, are revealed by variations in amino acids at the specificity determinants. These properties also result in similar interactions, such as hydrogen bonding (Fig. S5). This is expected, given the negative or neutral selection pressures across most sites in the DHp and Rec domains. With a few exceptions, such as DHp-aligned residues K245 (hydrophilic) and Y245 (hydrophobic depending on context and modification) and Rec D114 (hydrophilic), variants in all interacting residues in both domains are conserved in terms of hydrophobicity and hydrophilicity.

The observed high negative selection across the TCS sites indicates genetic constraints that shape their evolution in bacteria. This implies that mutations disrupting TCS gene function are generally harmful and likely eliminated by natural selection. As a result, the maintenance of new kinase-regulator pairs in the genome depends on intermediate steps between the original and new pairs being neutral or negative, allowing for the gradual evolution of new pairs without disrupting TCS system function. High negative selection pressures also indicate strong evolutionary constraints on these genes, potentially limiting bacterial adaptability to new environments or response to changing conditions. Considering the importance of TCS genes for bacterial survival and fitness, mutations disrupting their function are likely strongly selected against since they may negatively affect the ability of the bacteria to adapt to changing environmental conditions.

Amino acid coevolution and selection pressure analysis is fundamental for examining the specificity and molecular recognition in various HK-RR interactions(8, 61). From the coevolution analysis the interacting residues of the cognate HK and RR do not consistently exhibit a strong correlation (Fig. 7). We hypothesize that this may be attributed to the selection cutoff for the coevolving residues in the rec and DHp domain, which serves to minimize the coevolutionary signals between the interacting residues and reduce noise in the mutual information analysis. Our findings do indicate a strong coevolutionary relationship between certain adjacent residues and one or more of the interacting pairs. This may suggest that residues adjacent to the interacting residues might influence the interaction between the two proteins by indirectly affecting the interface through modulating the conformation, stability, or orientation of the interacting residues. The negative selection pressures on certain residues indicates that evolution preserves these crucial interactions. As such some TCS residues in the DHp and Rec exhibit coevolution patterns, while others do not, suggesting different evolutionary strategies.

Mutations in specificity residues require corresponding changes in their cognate regulators to maintain each pathway’s operation and avoid cross-talk as they diverged (13, 32). Identifying specific residues that contribute to this specificity is crucial, as TCS signaling pathways are not entirely insulated, as evidenced by the virulence regulation and global regulons. While this study focused on the overlapping regulons due to specificity at the phosphotransfer level DHp and Rec domain other determinants include interactions of the effector binding domain to specific motifs, receptor dimerization, or the response to specific environmental signals by the sensor kinase (8, 18).

In this study using multiple sequence alignments in conjunction with the cocrystal structure of the of the NarL DesK-DesR complex helped identify specific residues underlying phosphotransfer specificity between HKs and RRs. As genome sequence databases expand, covariation analysis has proven helpful in the absence of high-resolution structures of complexed cognate pairs. However, they cannot definitively identify the critical amino acid residues for specificity or reveal the degree to which each residue contributes to substrate selection. Experimental evidence such as Alanine scanning can provide further confirmation of specificity determinants in TCS.

In conclusion, an exhaustive exploration of signal fidelity determinants in TCS and their implications on virulence factor regulation in bacteria enriches our comprehension of bacterial evolution, adaptation, and the mechanisms governing their pathogenicity. This serves as a foundation for advancing synthetic biology through the manipulation of bacterial regulatory networks and rewiring of their modularity.

Considering the growing interest in targeting TCSs for anti-virulence strategies, a comprehensive understanding of their function, interactions, and specificity becomes paramount. This knowledge will not only provide insights into bacterial physiology but also inform more effective therapeutic interventions.

## ACKNOWLEDGEMENTS

Stephen Dela Ahator was funded by a grant from CANS.

